# Linked-Read Sequencing of Eight Falcons Reveals a Unique Genomic Architecture in Flux

**DOI:** 10.1101/2022.01.05.468466

**Authors:** Justin J.S. Wilcox, Barbara Arca-Ruibal, Jaime Samour, Victor Mateuta, Youssef Idaghdour, Stéphane Boissinot

## Abstract

Falcons are diverse birds of cultural and economic importance. They have undergone major lineage-specific chromosomal rearrangements, resulting in greatly-reduced chromosome counts relative to other birds. Here, we use 10X Genomics linked reads to provide new high-contiguity genomes for two gyrfalcons, a saker falcon, a lanner falcon, three subspecies of peregrine falcons, and the common kestrel. Assisted by a transcriptome sequenced from 22 gyrfalcon tissues, we annotate these genomes for a variety of genomic features, estimate historical demography, and then investigate genomic equilibrium in the context of falcon-specific chromosomal rearrangements. We find that falcon genomes are not in AT-GC equilibrium with a bias in mutations towards higher AT content; this bias is predominantly driven by—but not dependent on—hypermutability of CpG sites. Small indels and large structural variants were also biased towards insertions rather than deletions. Patterns of disequilibrium were linked to chromosomal rearrangements: falcons have lost GC content in regions that have fused to larger chromosomes from microchromosomes and gained GC content in regions of macrochromosomes that have translocated to microchromosomes. Inserted bases have accumulated on regions ancestrally belonging to microchromosomes, consistent with insertion-biased gene conversion. We also find an excess of interspersed repeats on regions of microchromosomes that have fused to macrochromosomes. Our results reveal that falcon genomes are in a state of flux. They further suggest that many of the key differences between microchromosomes and macrochromosomes are driven by differences in chromosome size, and indicate a clear role for recombination and biased-gene-conversion in determining genomic equilibrium.

**Significance:** Falcons are a particularly diverse and widespread genus of birds of particular cultural and economic importance. Falcons have also undergone recent large-scale chromosomal rearrangements to arrive at atypically low chromosome counts relative to other birds. We produced eight new high-quality falcon genomes to support general research on falcons, and we analyze these genomes to assess how chromosome loss has influenced specific aspects of genomic architecture.

## Introduction

Falcons (genus *Falco*) are among the most successful groups of birds, possessing an unparalleled geographic distribution (Gaston *et al.*, 2005; Gill and Donsker, 2019) and approximately 40 species, many resulting from several radiations in the last 3 million years (Fuchs *et al.*, 2015). This exceptional evolutionary and ecological diversity is coupled with a long and complex history of natural and cultural association with humans (Morgan and McGovern-Hoffman, 2008; Morin and Laroulandie, 2012), particularly falconry, which has transformed falcons into a living human heritage (UNESCO, 2016) and valuable commodities (Fleming *et al.*, 2011). Together, these unique natural, cultural, and economic aspects of falcons make them excellent subjects for genomic studies on a range of topics that span the gap of applied and basic science (Wilcox *et al.*, 2019).

The peculiar natural history of falcons is coupled with what is perhaps an even more peculiar genomic architecture. Through a series of interchromosomal rearrangements falcons have lost what was hundreds of millions of years of conserved synteny in theropods (O’Connor *et al.*, 2018a), and—at a chromosome count of 2N∼50—emerged with the lowest diploid chromosome number of any extant birds (O’Connor *et al.*, 2018b). This is primarily the result of a loss of microchromosomes <20MB in length (Axelsson *et al.*, 2005). Microchromosome loss is a fixed trait in falcons, and karyotypes within falcons are minimally divergent at 2N∼48-52, suggesting that microchromosomes losses occurred in the common ancestor of all falcons at least 7-10 MYA. While karyotypes have not been described for the closest relatives of falcons within the subfamily Falconinae, their next closest relatives, the Caracaras (family: Falconidae; subfamily: Polyborinae) possess typical avian karyotypes of 2N∼80-90 (Belterman and De Boer, 1990; Tagliarini *et al.*, 2007), suggesting that the chromosomal fusions in falcons occurred more recently than this divergence, i.e. within the last 15-20 million years (Fuchs *et al.*, 2015). While chromosomal fusions have been the subject of prior research (Joseph *et al.*, 2018), their evolutionary impact at the genomic level remains unexplored. Microchromosomes are distinct in their composition and behavior from larger chromosomes and are characterized by higher recombination rates, higher-mutation rate, higher gene density, more CpG-islands, and higher overall GC content (Axelsson *et al.*, 2005; Ellegren, 2010; Perry *et al.*, 2020; Schield *et al.*, 2020). Microchromosomes may also contain a higher density of structural variants (Völker *et al.*, 2010) and a lower density of repetitive elements (International Chicken Genome Sequencing Consortium, 2004). Debate continues as to whether the special characteristics of microchromosomes are a product of their size or a product of their sequence motifs. The fusions of microchromosomes in falcons provide an opportunity to assess this. Falcons have also been reported to have several other genomic peculiarities, including extensive nuclear mitochondrial DNA segments (NUMTs) relative to most other birds (Nacer *et al.*, 2017; Liang *et al.*, 2018), reduced TE abundance relative to other birds, and longer genic and intragenic lengths relative to other birds (Zhan *et al.*, 2014). Further investigation is necessary to determine links between these genomic peculiarities and the unusual chromosomal configurations of falcons.

Loss and fusions of microchomosomes could have profound effects on the genomic stability of falcons (Ellegren, 2010; Perry *et al.*, 2020). In particular, microchromosomes have been proposed as important for maintaining high-GC isochores and genomic AT-GC equilibrium (Duret *et al.*, 2006; Ellegren, 2010). Birds appear to be in or close to AT-GC equilibrium with high-GC isochores concentrated on microchromosomes (Costantini and Musto, 2017). Mammals for their part, lost their microchomosomes in a common ancestor more than 200 million years ago and are losing GC content from high-GC isochores (Duret *et al.* 2002). The Biased-Gene-Conversion Hypothesis (BGCH) proposes that high-GC isochores formed and are maintained by high-recombination rates, which favor conversion of GC-to-AT. More specifically, BGCH suggests that high-GC isochores in mammals originated from fused microchromosomes found in the common ancestor of extant amniotes (Duret *et al.* 2002), and are out of equilibrium due to subsequent reductions in recombination rate. Contrasting selectionist models have also been proposed for the origin of high-GC isochores and their ongoing decline in mammals (Costantini and Musto, 2017), and are based around the idea that a higher GC content confers higher chromosomal, mRNA, and protein stability at higher temperatures (Bernardi and Bernardi, 1986). These selectionist models have yet to find strong empirical support, despite a wide range of thermal systems on which to test these hypotheses (Videvall, 2018). In contrast the BGCH lacks supporting analyses in non-mammalian genomes.

Here, we sequence eight falcon genomes sampled from common falcons used in falconry: two gyrfalcons, a saker falcon, a lanner falcon, three subspecies of peregrine falcons, and the common kestrel (Table 1). These falcon species and subspecies represent diverse ecologies and are drawn from a cosmopolitan distribution. Saker falcons breed across central Eurasia and the Middle-East and take other birds and small vertebrates as prey. Gyrfalcons reside in the arctic of Eurasia and North America and take large birds as prey. Lanner falcons occur across Africa and the Mediterranean basin, as well as parts of the Middle-East and takes birds and some small mammals as prey. Peregrine falcons occur on every continent except Antarctica and typically take birds as prey: the subspecies used in this paper include the Eurasian peregrine (*F. p. peregrinus*), which occurs across temperate Eurasia; the black shaheen (*F. p. peregrinator*), which occurs in southern Asia; and, the Barbary falcon (*F. p. pelegrinoides*) which occurs in the Canary Islands, North Africa, and parts of the Middle-East. The common kestrel is much smaller and more distantly diverged from the other falcons included in our study. Like other kestrels, it preys primarily on small mammals (but can take birds), and occurs across Eurasia and northern Africa. Genomes are sequenced using 10X Genomics Chromium Linked-Reads sequencing, providing a first-ever genome for a lanner falcon, and greatly improved genomic assemblies for the saker falcon, and first-ever assemblies for the Barbary falcon and black shaheen subspecies of peregrines. We utilize the large phased scaffolds to examine compositional changes in falcon genomes and their partitioning between conserved and fused regions of the genome. We use alignments to related genomes to annotate the ancestral and current state of genomic regions, which allowed us to assess the BGCH by examining: 1) whether falcons are in AT-GC equilibrium, and 2) whether microchromosomes that have fused into larger chromosomes show a particular bias towards AT mutations.

**Table 1:**
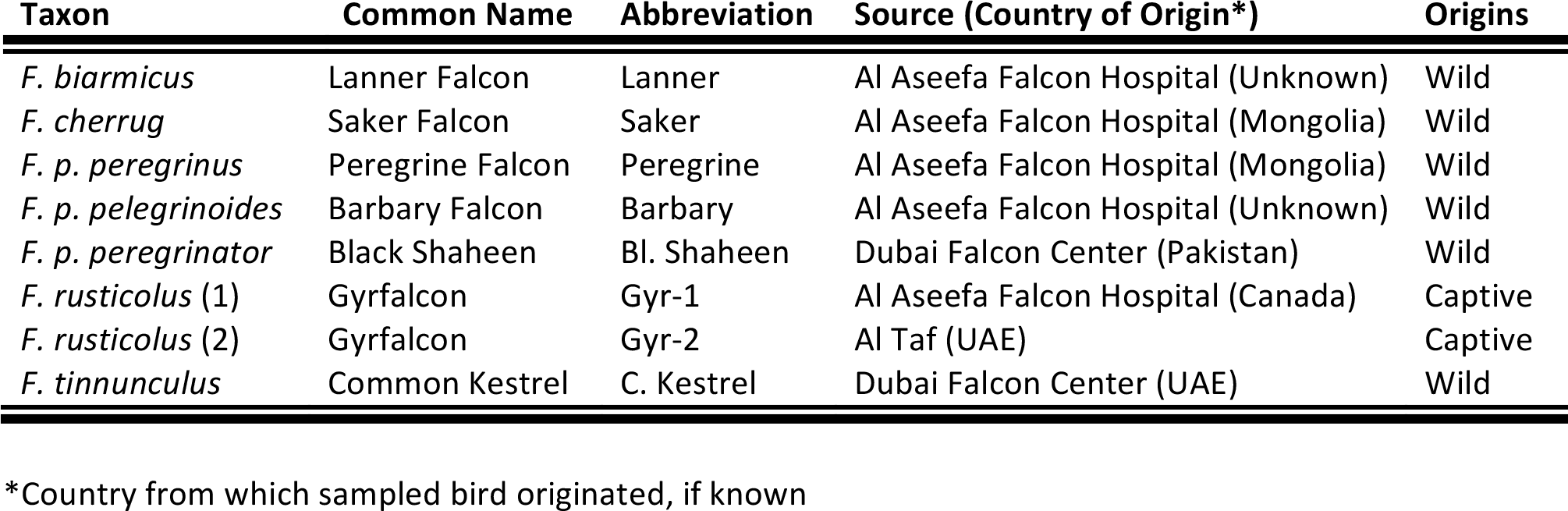
Falcon Sample Information

## Results

### Assembly Metrics

Reads were assessed for quality using FastQC. Average Pfred quality score per read was ∼36 across all genomes and we found no quality metrics of concern. We assessed for potential contamination, sequencing bias, and read bias using Kat. We found no evidence for contamination. Most genomes did however show a sequencing bias toward higher GC content, as indicated by a trend for higher GC content among more frequent k-mers (Supp. Fig. 1).

We sequenced 588 gigabases of DNA from eight falcon genomes using 10X Genomics Chromium linked-reads sequencing and assembled these using Supernova with assembly sizes ranging from 1.14 GB to 1.17 GB (Table 2). Scaffold counts ranged from 806 in the Black Shaheen to 2,041 in the common kestrel, with corresponding scaffold N50 values at a high of 40.62 MB and low of 10.58 MB. Raw coverage was greater than 50X and averaged 58X across all genomes, exceeding the 38X minimum coverage requirements and approximating the 56X optimum target recommended by 10X Genomics for assembly with Supernova. Effective coverage (i.e. with duplicates removed) ranged from 31.52 to 61.34X.

**Table 2:** Assembly Metrics

| Genome | Size(GB) | GC% | Scaffolds | Cov | N50 (MB) | BUSCOs: SC \| Dup \| F \| M* |
| --- | --- | --- | --- | --- | --- | --- |
| Lanner | 1.17 | 42.11 | 1591 | 39.22 | 28.94 | 94.4% \| 0.8% \| 1.0% \| 3.8% |
| Saker | 1.16 | 42.14 | 1152 | 43.67 | 29 | 95.1% \| 0.4% \| 0.9% \| 3.6% |
| Peregrine | 1.16 | 42.05 | 1136 | 40.72 | 25.88 | 94.8% \| 0.6% \| 1.0% \| 3.6% |
| Barbary | 1.16 | 42.04 | 1296 | 38.5 | 26.06 | 94.3% \| 0.3% \| 1.0% \| 4.4% |
| Bl. Shaheen | 1.16 | 42.02 | 806 | 61.34 | 40.62 | 93.8% \| 0.3% \| 1.0% \| 4.9% |
| Gyr-1 | 1.17 | 42.27 | 1518 | 45.59 | 20.6 | 95.1% \| 0.7% \| 1.0% \| 3.2% |
| Gyr-2 | 1.17 | 42.25 | 1699 | 43.29 | 17.12 | 95.2% \| 0.5% \| 1.2% \| 3.1% |
| C. Kestrel | 1.14 | 41.92 | 2041 | 37.52 | 10.58 | 91.1% \| 0.3% \| 1.3% \| 7.3% |

\*Out of 8338 BUSCOs: SC=Complete Single-Copy | Dup=Complete Duplicated | F=Fragmented |
M=Missing

Genomic completeness was assessed using BUSCO to search for single-copy avian orthologs with the “aves_odb10” database containing 8338 universal single-copy avian orthologous genes (Table 2). The proportion of BUSCO genes detected as single-copy and complete ranged from 91.1% for the common kestrel to 95.2% for Gyr-2 with a median of 94.6% across all assemblies. The proportion of BUSCOs that could not be detected (i.e. “missing” BUSCOs) ranged from 3.1% of the database in one of the gyr falcon (Gyr-2) to 7.3% of the database in the kestrel; across all assemblies a median of 3.7% of BUSCOs were not detected.

The presence of scaffolds belonging to the Z and W sex chromosomes were confirmed in all falcon genomes using HMMER. Two close proximity hits to sex chromosome specific Z (CHD-Z) and W (CHD-W) Chromosome Helicase genes (Fridolfsson *et al.*, 1999) were found on separate scaffolds in all genomes. This search also produced a hit to a short segment of a predicted CHD-2 gene on a third scaffold of all genomes. No other hits were found to our HMMER model in any genomes.

Single Nucleotide Variants (SNVs) and small indel variants were called using three methods: long-ranger, MUMmer alignment to the common kestrel genome, and MUMmer alignment of phased diploid genomes to one another. The number of heterozygous sites among small variants was comparable across methods (Supp. Fig. 2). Heterozygous variants ranged from 623,505 SNVs/ 139,591 indels in Gyr-1 to 4,578,621 SNVs / 627,414 indels in the common kestrel using the self-alignment; and 820,996 SNVs / 155,477 indels in Gyr-1 and 4,952,485 SNVs / 705,344 indels in the common kestrel using Longranger. We detect ∼86% more heterozygous SNVs in the saker falcon and ∼42% more SNVs in the peregrine than reported in previously published genomes for these species (Zhan *et al.*, 2013). We detect ∼4% fewer SNVs than previously reported for the common kestrel (Cho *et al.*, 2019). The average distance between heterozygous sites varied greatly from 0.250KB in the common kestrel to 1.75KB in Gyr-1 (Supp. Table 1).

Structural variants were annotated from MUMmer alignments to both the kestrel genome and phased haplotypes of each individual genome. Structural variants private to each diploid genome varied from approximately 1000 to 1500 in number (Supp. Fig. 3A). By count, unique structural variants were dominated by inversions and segmental duplications. Numbers of insertions and deletions varied between genomes but these accounted for only a small portion of structural variants in all genomes. Unique segmental deletions were consistent in number across genomes and ranged in count from 28 in the kestrel to 51 in the saker falcon. Unique structural variants varied in size distributions across types, and as such, counts did not reflect the differences in base pairs attributed to each type of structural variant (Supp. Fig. 3B). Base pairs of structural variation were generally dominated by inversions, which ranged in diploid counts from 1,123,579 bp in the kestrel to 4,720,786 bp in Gyr-1. Unique inserted base pairs were extremely variable in number, and ranged from diploid counts of 1,102,918 bp in the Gyr-1, to 42,404,444 bp in the Eurasian peregrine and 78,289,714 in the Black Shaheen, with between 1.5 and 3.5 million unique inserted bases in the other falcons.

We performed a *de novo* repeat annotation by building a database of repeat subfamilies with RepeatModeler 2 and searching genomes for these with RepeatMasker using rmBLAST. The proportion of repeat elements annotated in each genome ranged from 6.38% in the Kestrel to 6.61% in Gyr-1. Repeat element composition was dominated by LINEs of the Chicken Repeat 1 (CR1) family with a very small number of L2 and Penelope elements detected. A small number of Long Terminal Repeat (LTR) and ‘cut-and-paste’ DNA elements were also detected. The divergence curves showed the same pattern for all samples with the vast majority of repeats being more than 10% divergent from consensus sequences and a particularly strong wave of amplification between 10% and 15% divergence and a somewhat older wave of amplification at 25-30% divergence for several repeat families (Figure 1). To approximate the times of these amplifications we used the same *de novo* libraries from falcons to annotate repeats in birds from other clades of Eufalconimorphae (the Swainson’s Thrush and Kakapo) and an outgroup within Australaves (the red-legged Seriema). The annotation of repeats in these other genomes, suggest that both waves of amplification were already present at the time of the divergence of the Eufalconimorphae from its common ancestor with seriemas ∼60MYA (Prum *et al.*, 2015). Near identical repeat landscapes between the falcons and red-legged seriema—and similar patterns in the kakapo—suggest minimal activity for the annotated repeat subfamilies in these lineages. This finding is supported by a lack of intact ORFs belonging to repeats in falcons. Across all genomes we identify 14,532-15,558 ORFs with matches to active LINE elements, but all of these had premature stop codons. We identify 10-28 ORFs belonging to LTR elements across all genomes, again all with premature stop codons. No ORFs belonging to DNA transposons were identified. Finally, we find no unique insertion variants corresponding to annotated repeats that are greater than 300bp in any genome. Interspersed repeats were not evenly distributed across the genome. Base-pairs arising from transposons and retrotransponsons were found to be significantly enriched on 100 KB windows of the Z chromosome relative to autosomes (p< 1e-6).Fusions of chromosomes could result in interstitial telomere which can be detected in the simple repeats annotated by RepeatMasker. In light of recent chromosomal fusions in falcons we searched for simple repeats that matched either the vertebrate forward (TTAGGG) or reverse (CCCTAA) telomere sequences and that were not located on the beginning or end of a scaffold. We found between 16 and 26 simple repeats corresponding to interstitial telomeres within each diploid falcon genome.

**Figure 1:**
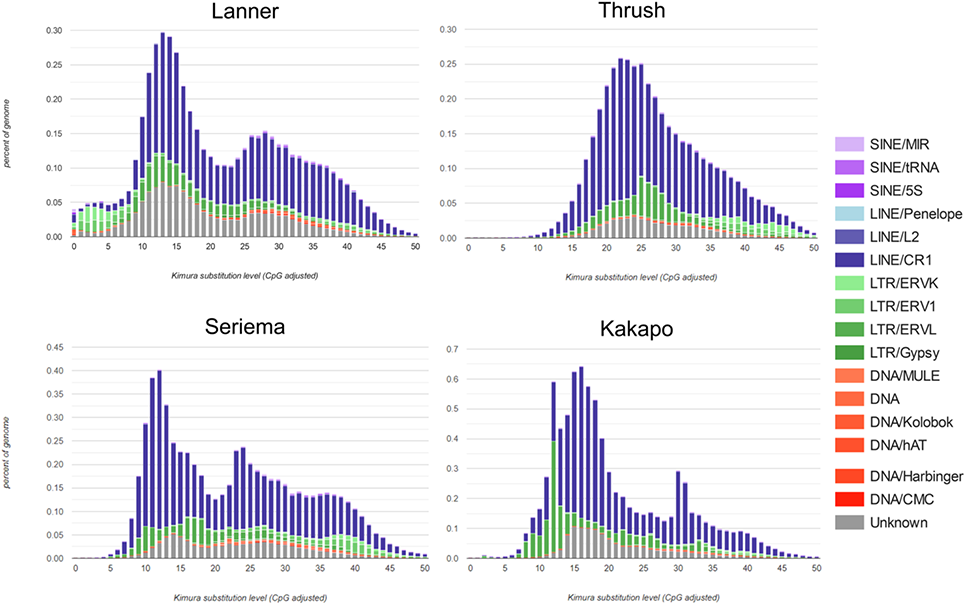
CpG adjusted divergence of repetitive elements from consensus sequences for the lanner falcon (representative of other falcons) and three other from within Eufalconimorphae sampled from NCBI RefSeq and Genbank: the Swainson’s thrush; the kakapo; and, red-legged seriema. Repetitive elements are color coded by type and family. All genomes were annotated with dereplicated *de novo* repeat libraries compiled and dereplicated from the eight falcon genomes.

We annotated 100 KB (Supp. Figure 4) and 1MB (Supp. Figure 5) windows to assess distributions of GC content throughout falcon genomes in the context of isochores (Costantini and Musto, 2017). Falcon genomes showed similar patterns of GC distribution to one another and other birds (Figure 2). Avian genomes most closely resembled the human, but with a slight shift toward higher GC content.

**Figure 2:**
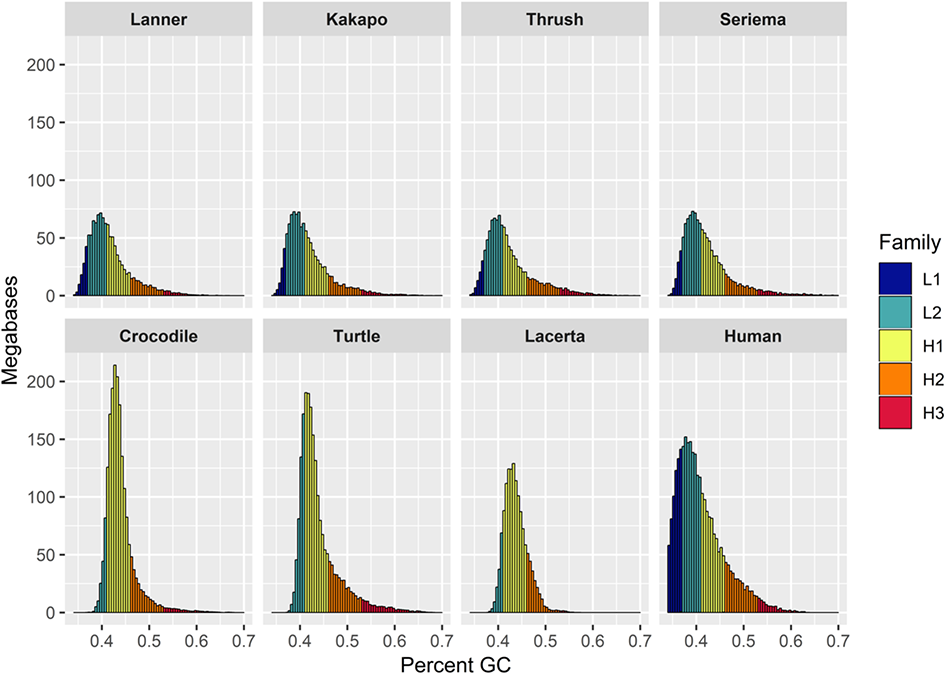
Distributions of GC content across 100 KB windows for the representative lanner falcon and other Eufalconimorphae and amniote lineages sampled from NCBI RefSeq and Genbank: the kakapo; the Swainson’s thrush; the red-legged seriema; the saltwater crocodile (*Crocodylus porosus*); the green sea turtle (*Chelonia mydas*); the sand lizard (*Lacerta agilis*); and human (*Homo sapiens*). Isochore families are delineated by color.

### Evolutionary History

A Maximum-Likelihood phylogeny was constructed in RAxML using all 15,426,039 variable SNV sites derived from the MUMmer alignments to the haploid kestrel genome (Figure 3). The subgenus *Hierofalco* (i.e. the lanner falcon, saker falcon and gyrfalcon) and peregrine falcons fell out into clearly defined clades. Within *Hierofalco*, the two Gyr genomes grouped together along with the saker and lanner diverging ancestrally to these. All groupings had 100% bootstrap support. The tree was scaled to the divergence of old-world kestrels and large falcons at the Tortonian-Messinian Junction, 7.246 MYA, in the late Miocene (Boev, 2011). It suggests a split between the peregrine falcons and *Hierofalco* of approximately 588 KYA with a split between the lanner and saker falcon of approximately 304 KYA. The saker and gyrfalcon and the peregrine subspecies all respectively coalesced within the last 150 KYA.

**Figure 3:**
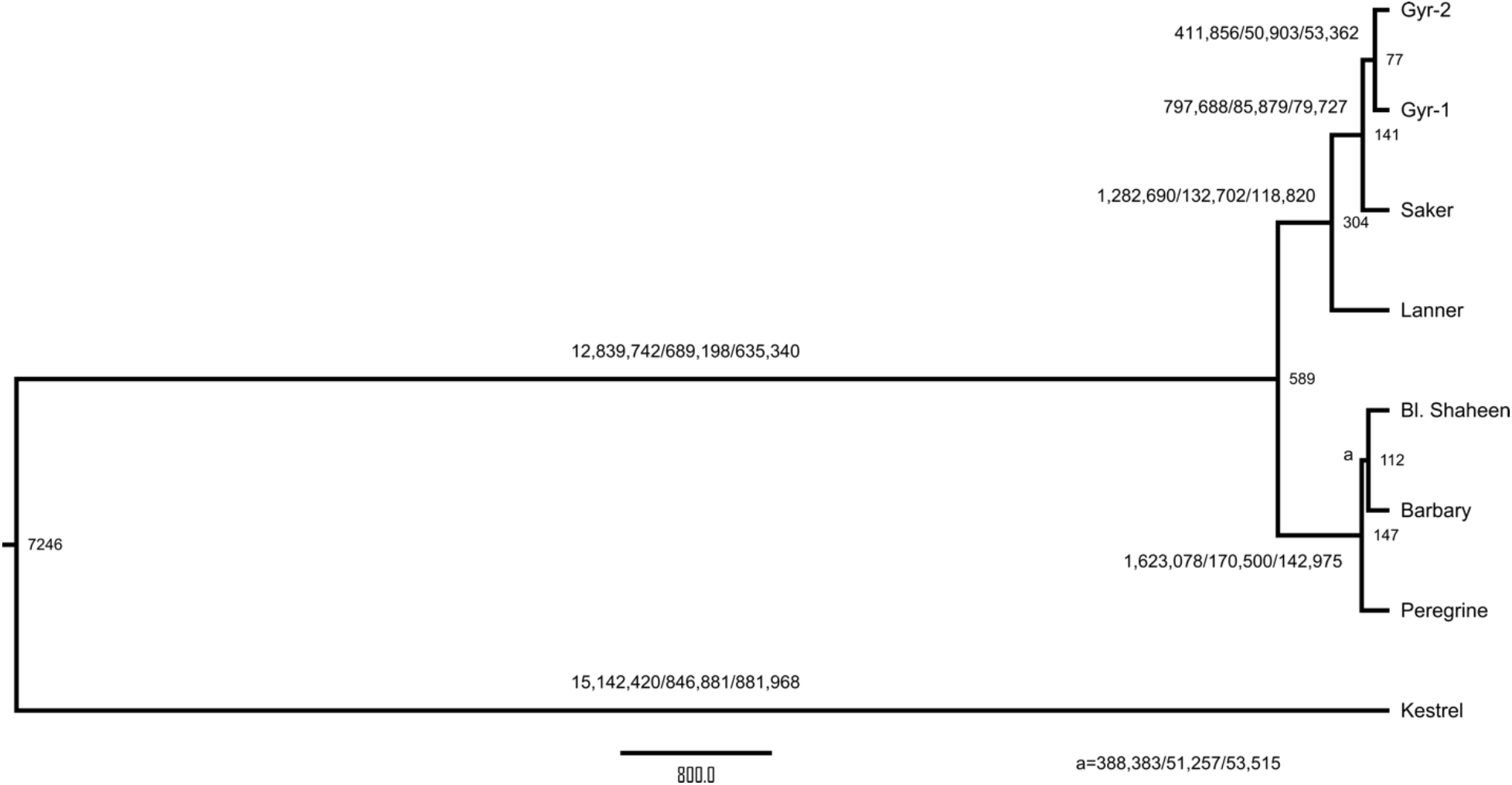
Maximum-likelihood tree of the eight diploid falcon genomes time-scaled by the divergence of the common kestrel from other falcons. Tree was built using all variable SNV sites detected from alignment to the kestrel genome. Branch labels denote bootstrap values based on 1000 bootstraps. Scale bar denotes substitution rate. Branch labels denote the branch specific number of unique shared small variants <50 bp: SNVs/Deletions/Insertions.

PSMCs were performed to reconstruct the demographic history contributing to each genome (Figure 4). Analyses of the peregrine falcons and subgenus *Hierofalco* genomes suggest larger effective population sizes ∼4-8 MYA followed by substantially reduced population sizes (Supp. Fig. 6). The saker falcon, lanner falcon and peregrine falcons all show evidence for a recent population expansion and more recent population contraction within the last few hundred thousand years. The expansion and contraction were very pronounced in the lanner falcon and Eurasian peregrine falcon, but very muted in the other falcons. The common kestrel followed a trend that was reversed from the large falcons, with a lower ancestral population size and several recent expansions. Scaling suggests that these expansions began approximately one million years ago, with the most recent waves of expansion occurring within the last two-hundred-thousand years.

**Figure 4:**
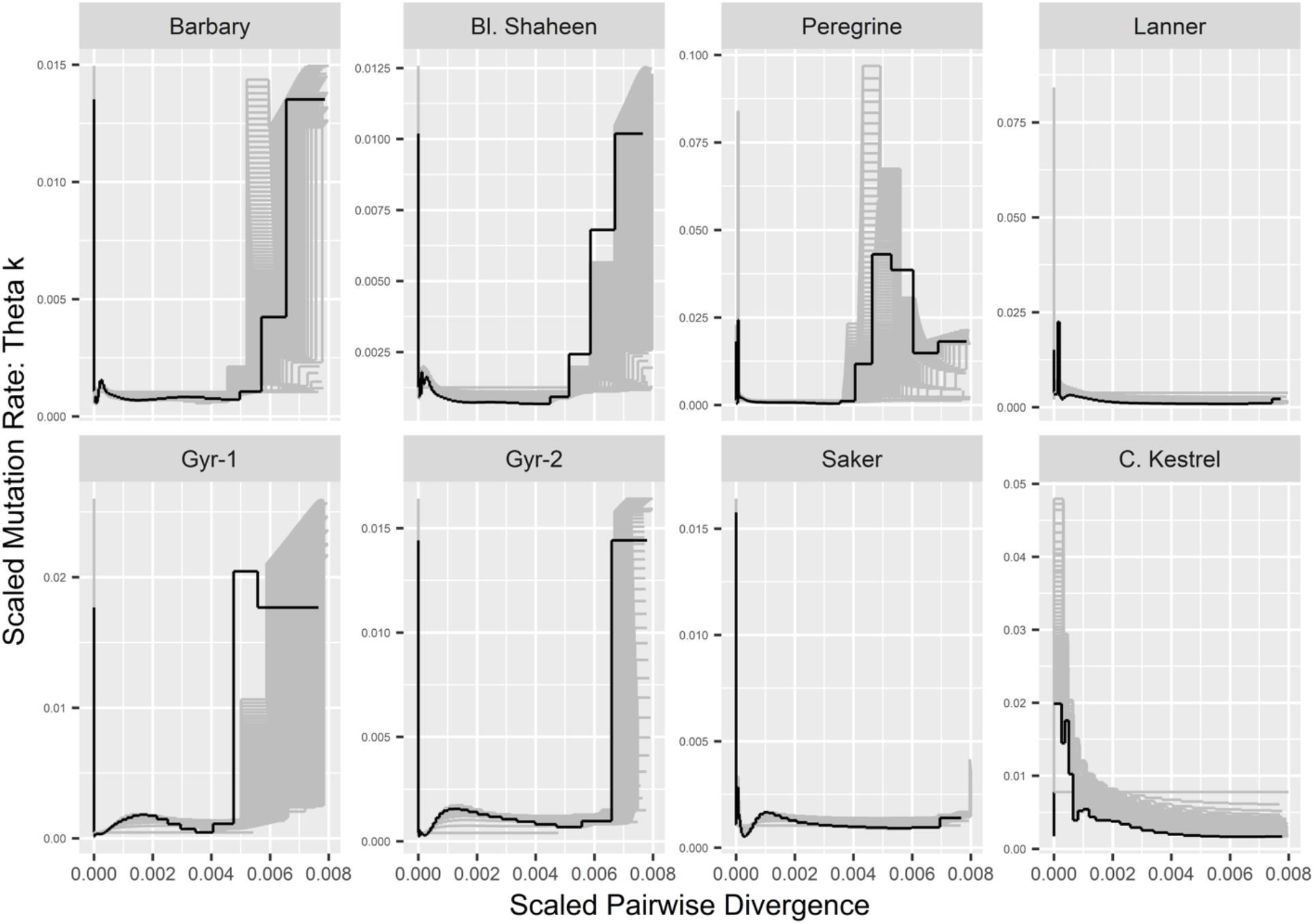
PSMC results of demographic trends for each falcon genome. Time is indicated on the X-axis and is represented by scaled pairwise divergence, with higher values indicating more ancestral time points. Population size is indicated on the Y-Axis as scaled population mutation rate, with higher values indicating larger effective population sizes. Bootstraps were performed in 100 replicates and are shown in grey lines.

### NUMTs evolution

We annotated 900 NUMTs accounting for 992,144 base pairs across the eight falcon genomes of which 316 were unique to specific genomes, totaling 368,339 base pairs. We annotated 102-116 total NUMTs in each diploid genome, consisting of 105,585-150,107 total bases (Supp. Table 2). The number of heterozygous NUMT insertions (those occurring on only one haplotype) varied widely, ranging from 13.2% in Gyr-1 to 55.2% in the kestrel with an average of 27.9% across all genomes; the majority of NUMTs were found on both haplotypes of all genomes except the common kestrel. The proportion of NUMTs with syntenous insertions in other genomes was relatively invariant in the large falcons with a range of 55.5%-72.9% of NUMTs and an average of 61.5%. The exception to this was again the kestrel, in which only 20.7% of NUMTs had shared synteny with other genomes. We conducted a phylogenetic analysis of NUMTs by clustering them at 80% similarity, aligning clusters with mitochondrial genomes of taxa from across avian evolution and, constructing maximum-likelihood trees from each cluster. NUMTs formed 48 clusters at 80% similarity. Across all clusters trees were generally grouped by synteny (Generalized Linear Models: weighted-mean-of-R^2^=0.729; weighted-standard-deviation-of-R^2^=0.051). The length of NUMTs was also overwhelmingly explained by synteny (Generalized Linear Models: weighted-mean-of-R^2^=0.81; weighted-standard-deviation-of-R^2^=0.036). Phylogenetic analysis suggests that the NUMTs found in falcons have been inserted from across the evolution of Aves, with a small number that have inserted before the divergence between the ancestors of Neoaves and Palaeognathae ∼72 MYA (Figure 5). Out of 900 NUMTs, we were able to determine the evolutionary origin of 571 using a phylogenetic analysis. The majority (N=454) of these NUMTs have inserted since the divergence of Falconidae from other Neoaves in the late Paleocene (∼56 MYA) with large burst of NUMT insertions predating divergence of the ancestors of Falconinae (i.e. the “true” falcons and kestrels) and Polyborinae (i.e. the caracaras) from Herptherinae (i.e. the laughing and forest “falcons”), and another burst of insertions predating the divergence of Polyborinae from Falconinae. In total 158 NUMTs clustered with or within *Falco* with the majority (N=99) clustering most closely to a kestrel mitogenome.

**Figure 5:**
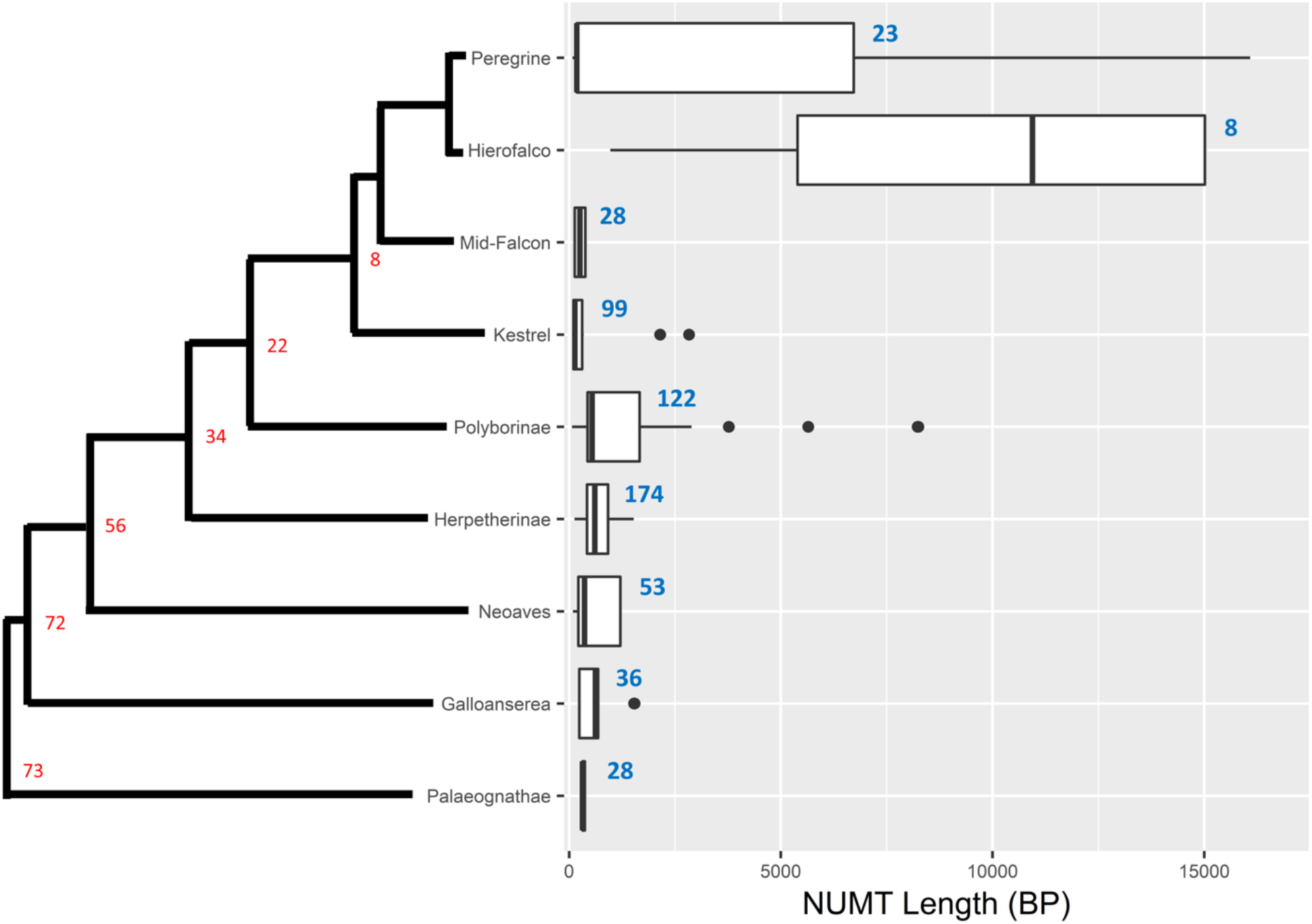
Basepair length (x-axis) distributions and number (labeled on graph in blue) of NUMTs estimated to have inserted in the common ancestor of falcons and the indicated lineages of Aves (y-axis) based on maximum-likelihood analysis. Boxplots show 1^st^ and 3^rd^ quartiles with median; lines denoted 1.5X interquartile range, with outliers outside this shown as circular dots. The tree to the left of the Y-Axis shows the evolutionary history based on an ultrametric Bayesian analysis of mitogenomes. Estimated divergence times are reported from literature (Prum *et al.*, 2015) on nodes in red.

We examined the propensity of NUMTs to insert into microchromosomes versus larger chromosomes by comparing the proportion of NUMTs inserted into 100 KB windows of each chromosome type with the proportion of 100 KB windows annotated as belonging to each chromosome type. We found no NUMTs that had inserted onto microchromosomes in falcons, constituting a significant propensity for NUMTs to insert onto larger chromosomes, although this finding did not constitute a significant bias after controlling for multiple tests (Binomial Exact Test: p=0.002478, N=158; Odds Ratio 95% Confidence Interval: 0.0-0.02307; Null Probability: 0.02755). We also compared the proportion of NUMTs inserted into 100 KB windows of each chromosome type before the divergence of falcons with the proportion of 100 KB windows estimated to have ancestrally belonged to each chromosome type. We again found no NUMT insertions onto ancestral microchromosomes, signifying a significant propensity for NUMTs to insert onto larger chromosomes (Binomial Exact Test: p<1e-6, N=413; Odds Ratio 95% Confidence Interval: 0-0.00889; Null Probability: 0.1016). We find that newer NUMT insertions are significantly longer than older NUMT insertions using two independent approaches: we find that the length of NUMTs are negatively correlated with the distance of NUMTs from the mitochondrial genome of the falcon in which they were detected (Generalized-Linear-Model-with-Poisson-Error-Distribution: p<1e-6; parameters estimate: −0.381 base pairs per substation per base pair); insertions within *Falco* were also significantly longer than insertions in the vicinity of divergence from other Falconidae, other Neoaves, and Galloanserea (Bonferoni-Corrected-Pairwise-Wilcoxon-Rank-Tests: p<0.0001).

### Genomic Stability and Disequilibrium

To assess whether falcon genomes were in AT-GC equilibrium, we analyzed SNVs private to each falcon genome (Supp. Fig. 7A). SNVs were called using MUMmer alignments to the kestrel genome. SNV counts were weighted such that those that were fixed on both haplotypes of a falcon genome were counted twice and those that were heterozygous were counted once. Focusing on private mutations effectively used a maximum-parsimony approach for assigning ancestral state, and avoided pseudoreplication of the same mutation events across multiple genomes. It should also have the advantage of biasing our analyses toward more recent mutations. Focusing on SNVs that are unique to each genome may also bias our analysis slightly in the direction of sequencing error. We note that Novaseq base calling errors have a slight bias (<20% higher likelihood of error) toward higher GC content (Ma *et al.*, 2019). This may conceal some loss of GC content in our analysis, but overall low error rates (<1E-4) in SNV calls should make the impact of miscalls on our analysis negligible. The odds ratios of sites shifting from A or T to G or C were compared. Analyses were performed with and without CpG sites included; analyses without CpG excluded any sites that contained CpG in any of the eight genomes. Mutations were out of AT-GC equilibrium, and biased toward increasing AT content with w/CpG sites included (Odds-Ratio=1.545; Chi-Squared= 327,100; p< 2.2×10^-16^) and w/o CpG (Odds-Ratio=1.103; Chi-Squared= 10,100; p< 2.2×10^-16^). Universally higher odds-ratios were observed for analyses with CpG sites (Sup. Fig. 7B).

We assessed insertion and deletion variants to determine if genome size was in equilibrium using only insertion and deletion sites that were private to each genome, in a manner that paralleled our handling of mutational biases in SNVs above. Among small variants, insertions were significantly longer than deletions for all peregrine falcons and *Hierofalco* genomes (Welch-t= 39.395; p < 1e-6), but shorter than deletions in the kestrel genome (Welch-t= 39.395; p-value < 2e-16; Supp. Fig. 8). Binomial exact tests found that insertions were also significantly more common than deletions for all genomes except the kestrel (< 1e-6), in which significantly more insertions were found (p< 1e-6). Similar results were found for structural variants: insertion variants were both significantly more numerous (p=0.00152) and had significantly more base pairs than deletion variants (p< 1e-6). Segmental duplications were significantly more numerous than segmental deletions (p< 1e-6), but no differences were found in their average size.

Finally, we assessed genomic equilibrium in the context of isochores and ancestral and current chromosomal states, as delineated by 100 KB windows (Costantini and Musto, 2017). We defined ancestral chromosomes types of windows based on their universally conserved alignments to chromosomes of this type in select chromosome-scale assemblies sampled from the other lineages of Australaves (i.e. seriemas, parrots, and song birds), using the: red-legged seriema (Cariamiformes), kakapo (Psittaciformes), and Swainson’s thrush (Passeriformes). We began by testing for differences in GC content between 100 KB windows based on differences in ancestral and current chromosomal type—microchomosomes and large chromosomes (log-transformed ANOVA: F= 5485.723; p<1e-6; Figure 6A; supp. Fig. 9). Windows ancestrally belonging to microchromosomes were associated with higher GC content relative to regions that were conserved on larger chromosomes (p<1e-6). Moreover, windows that were ancestrally found on microchromosomes but now fused into larger chromosomes had lower GC content than those conserved on microchromosomes (p<1e-6), indicating that microchromosomes fused onto larger chromosomes have lost GC content. Windows ancestrally located on large chromosomes but now on microchromosomes, likewise had higher GC content than those conserved on large chromosomes (p<1e-6), indicating convergence towards higher GC content. A similar pattern was found for the percent CpG sites with windows that had fused from microchromosomes into larger chromosomes possessing a higher proportion of CpG relative to those conserved on large chromosomes and a lower level of CpG than those conserved on microchromosomes (log-transformed ANOVA: F= 252.95; p<1e-6; Figure 6B; Figure 6Bl; supp. Fig. 10). When we control for GC content, we observe the same pattern (log-transformed ANOVA: F= 28.37; p<1e-6; supp. Fig. 11), except that windows ancestrally on microchromosomes and fused onto large chromosomes are found to be depleted in CpG relative windows that are conserved on large chromosomes, suggesting that microchromsomes that have fused onto larger chromosomes have depleted in CpG more quickly than GC.

**Figure 6:**
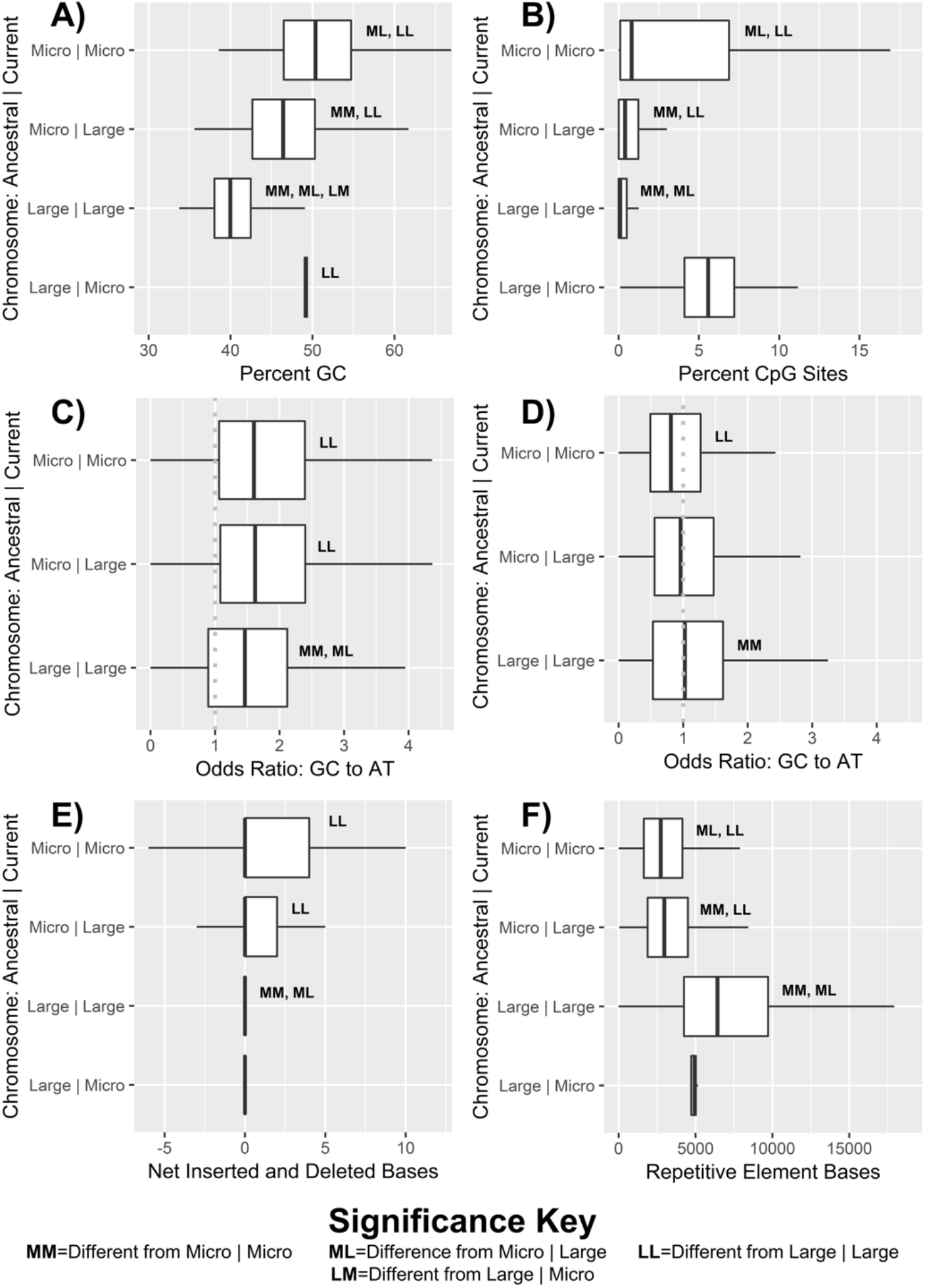
Boxplots of genomic features by Ancestral | Current chromosomal state of 100 KB windows across the eight falcon genomes for: A) Percent GC content; B) Percent CpG sites; C) Odds ratios of AT to GC mutation with CpG sites included; D) Odds ratios of AT to GC mutation with CpG sites excluded; E) Net inserted and deleted bases; F) Total number of bases annotated by as belonging to TEs. Significant differences are denoted with letters according to descriptions in the bottom key.

The AT-GC equilibria of private mutations also differed based on current and ancestral chromosomes state (log-transformed ANOVA: F=76.271; p<1e-6; Figure 6C; supp. Figure 12). Windows that were ancestrally found on microchromomes were found, regardless of current chromosome type, to be losing GC content at higher rates than regions that were conserved on large chromosomes (p<1e-6). These results were largely driven by a trend toward AT biased mutation in regions with high GC content (p-value < 2e-6). After regressing out the effect of GC content, a significant effect of chromosome state was still observed (log-transformed ANOVA: F= 8.930; p=0.000133; supp. Figure 13). Windows ancestrally belonging to microchromosomes but fused into larger chromosomes had a significantly (p= 0.0004) higher mutational bias toward AT than those windows with conserved status on large chromosomes, indicating that regions of microchromosomes fused onto large chromosomes are still losing GC content at rates higher than expected for their GC content. In contrast, regions that were conserved on microchromosomes and macrochromosomes showed no differences in GC loss when controlling for GC content, indicating that conserved regions are losing GC content at rates consistent with their GC content. When CpG sites are removed weaker differences are observed between regions based on chromosome state (log-transformed ANOVA: F= 13.540; p<2e-6; Figure 6D supp. Figure 14), and these are characterized by a reverse in trends with regions conserved on microchromosomes showing significantly lower bias towards AT content than regions conserved on larger chromosomes (p=3e-6). These differences were also largely explained by differences in GC content, which remained strongly correlated to mutational bias (p<1e-6). No effect of chromosomal state was found when controlling for GC content in the absence of CpG sites (log-transformed ANOVA: F= 2.28; p=0.132; supp. Figure 15), suggesting that recent mutational biases between former microchromosomes regions fused to macrochromosomes are driven by loss of CpG sites.

To determine if observed biases in inserted base-pairs were related to chromosomal fusions, we assess net inserted or deleted base pairs across 100 KB windows in the context of current and ancestral chromosomal state (log-transformed ANOVA: F= 9.768; p<2e-06; Figure 6E; supp. Figure 16). Regions formerly on microchromosomes and fused to large chromosomes had more inserted bases than windows conserved on large chromosomes (p=0.000064), as did windows with conserved positions on microchromosomes (p=0.00125). But, no significant difference was observed between windows formerly on microchromosomes that had fused with larger chromosomes and those that had not. Model selection criteria did not indicate GC content as important to this model so it was not included. The total number of small indels was, however, strongly correlated to GC content (p<1e-6). After controlling for GC content we found that windows ancestrally belonging to microchromosomes had significantly more indels than those that belonged to large chromosomes (p<1e-6). While there was a trend toward more indels on windows that had remained on microchromosomes relative to windows that had fused to large chromosomes from microchromosomes, this trend was non-significant after controlling for multiple tests (p=0.002). Counts of unique large structural variants were generally independent of both current and ancestral chromosome state, but were significantly depleted on microchromosomes fused to large chromosomes relative to large chromosomes after controlling for GC content (p=0.00009).

Finally, we looked at changes in chromosome state and base pairs annotated as belonging to transposable elements (TEs) across windows (log-transformed ANOVA: F= 1566; p<1e-06; Figure 6F; supp. Figure 17). Microchromosomes had fewer repeats than macrochromosomes and regions formerly belonging to microchromosomes but fused to large chromosomes had intermediate numbers of repetitive bases (p<1e=6). Higher GC content was found to be strongly negatively correlated with TE base content (p<1e-6), and the effect of GC content was regressed out (log-transformed ANOVA: F= 300.5; p<2e-06; Figure 6F; supp. Figure 18). After controlling for GC content, we found that windows ancestrally belonging to microchromosomes have fewer repetitive bases than those conserved on larger chromosomes (p-value< 1e-6). However, windows formerly belonging to microchromosomes but fused onto large chromosomes were nonetheless depleted in repetitive bases relative to those conserved on microchromosomes (p-value=0.0005). In light of a confirmed loss of GC content, this reversal of relationships when accounting for GC content, suggests a more rapid loss of bases associated with TEs than GC content on regions of microchromosomes that have fused to large chromosomes.

## Discussion

### Overview

We sequence the genomes of eight falcons using 10X Genomics Linked-Reads and produce highly-contiguous scaffold level assemblies that exceed *de novo* assembly sizes for all previously sequenced falcon species other than the gyrfalcon and the common kestrel, and at assembly sizes comparable to those reported for all previously sequenced falcons. We find genetic variation, particularly SNVs, to be highly variable across species with an ∼7X greater distance between heterozygous sites in gyrfalcons relative to the common kestrel. Using the Trinnotate pipeline (Bryant, 2017) we annotate a number of genes comparable to those found in previously published falcon genomes. We find distributions of GC content and isochore families similar to those reported in other birds. Bird genomes resembled mammalian genomes most closely, but were shifted toward higher GC content as reported in previous studies (Constantini and Musto, 2017). We annotate ∼6.5% of the falcon genomes as belonging to repetitive elements, a number comparable to reference-based analyses of saker and peregrine falcon genomes (Zhan *et al.*, 2013) and higher than a previous *de novo* annotation of the peregrine genome (Zhang *et al.*, 2014). We also find broad evidence for a lack of repeat activity in falcons. This finding is consistent with previous reports—using standardized methodology—that show falcons to be in the lowest quartile for repeat content in birds (Zhang *et al.*, 2014). We also find simple repeats corresponding to interstitial telomeres. While interstitial telomeres are known to be common in ratites (Palaeognathae), land fowl (Galloanserae), and waterfowl (Order: Anseriformes), they are not known to be common in the clade consisting of falcons, parrots, seriema and songbirds (Nanda *et al.*, 2002).

### Demography

We conduct a PSMC to assess changes in the population sizes of falcons over time. Our findings suggest that the population sizes of large falcons have declined since the time of their shared common ancestor. Based on scaling and previous estimates of divergence times between species we would estimate that these larger population sizes occurred in a shared common ancestor or common ancestors of large falcons 2-5 MYA (Fuchs *et al.*, 2015). Falcons have undergone a series of recent radiations on this time scale giving rise to the majority of extant falcon species. These large effective ancestral population sizes and subsequent reductions could reflect population subdivision as reproductive barriers arose during these speciation events (Ottenburghs *et al.*, 2017). The time scale of larger population sizes in large falcons would likely date back to the Pliocene with reductions potentially occurring during the transition from the Pliocene to the Pleistocene. Many avian populations display similar reductions in population size during this period (Nadachowska-Brzyska *et al.* 2015). Glaciation is the most obvious explanation for these reductions and glaciation is also a likely explanation for the reductions in population sizes observed in large falcons. Notably, common kestrel populations do not follow the same trends as those of the large falcons. We note that the ancestors of common kestrels did not undergo major radiations as recently as the large falcons (Fuchs *et al.*, 2015). Common kestrels would seem to have undergone a relatively recent and major population expansion, likely within the last few hundred thousand years. Similar major expansions were noted in the Eurasian peregrine and lanner falcon, although these were followed by more recent and rapid contractions. Glacial recession may have played a role in these events (Nadachowska-Brzyska *et al.* 2015), and as in the case of the larger falcons, introgressions cannot be ruled out (Hawks, 2017). Ultimately, studies on more falcon species and more genomes may provide additional insights into the underlying reasons for falcon population fluctuations in the more recent and distant past (Spence *et al.*, 2018; Gu *et al.*, 2021).

### NUMTs

We also annotate several dozen NUMTs in each falcon genome. While we annotate somewhat more NUMTs, these findings are consistent with previous reports from falcons (Nacer and do Amaral, 2017;), and are higher than typical of other birds (Liang *et al.*, 2018). Most but not all NUMTs shared synteny with NUMTs in other genomes. Phylogenetic analysis also showed that all NUMTs grouped most closely with syntenous NUMTs when these occurred. These findings suggest that primary insertion is the overwhelming source of NUMTs in falcons, with secondary duplication playing only a minor role. We find strong evidence that the NUMTs found in falcons have been accumulating across avian evolution, with the earliest likely predating the split between Ratites and other birds. However, most NUMTs inserted within the early evolution of Falconidae. This could suggest something particular to Falconidae that favored NUMT insertion, but could also suggest slow but continual decay of older NUMTs within the genomes of falcons and their ancestors. Genomes from other Falconidae will undoubtedly help to clarify these possibilities when they become available. Insertions within falcons are also ongoing. We find support for the hypothesis of an ongoing reduction in length of NUMT insertion over time, possibly favored by selection for smaller genomes in birds (Kapusta *at al*., 2017). We also propose—based on our findings of large insertion in large falcons but not the common kestrel—that NUMT insertion may be more common and larger in species with smaller effective population sizes, as smaller population sizes should increase the likelihood of genomic fixation for novel insertions due to drift and decrease the efficiency of selection against such insertions. This could explain higher numbers of NUMTs in the carnivorous Falconidae lineage, but this is a hypothesis better analyzed across a large number of species. NUMTs were biased against insertion on microchromosomes, and as such microchromosome loss could favor NUMT insertion. However, given the large accumulation of NUMTs that appear to predate the divergence of falcons from caracaras (and therefore the loss of microchromosomes in falcons), higher numbers of NUMTs within falcons cannot be generally attributed to their unique chromosomal rearrangements.

### Chromosomal Fusions and a Genome in Flux

We find that falcon genomes are out of AT-GC equilibrium and that they are losing GC content. Loss of GC content appears to be genome-wide, but, as in mammals (Duret et al. 2002), it is most pronounced in high-GC regions. Moreover, we find evidence that loss of GC content is linked to the loss of microchromosomes in falcons: regions of microchromosomes that have fused to larger chromosomes are depleted in GC content relative to regions that have remained on microchromosomes; and, regions of microchromosomes that have fused to larger chromosomes are losing GC content at a higher rate than typical of the larger chromosomes on which they now occur, even when controlling for higher GC content. Conversely, regions of larger chromosomes that have merged to microchromosomes appear to have gained GC content, although these sites were too rare to assess recent mutational biases. Historic loss of GC content is paralleled by a loss of CpG islands, although the ongoing loss of GC content is driven largely but not exclusively by the loss of CpG sites.

Overall, our findings provide support for the Biased-Gene-Conversion Hypothesis (Duret *et al.*, 2006) as a driver of genomic mutational disequilibrium in falcons. Microchromosomes that have fused to larger chromosomes are presumably subject to reduced recombination rates as a result of a greater number of base pairs over which recombination events may occur (Ellegren, 2010; Perry *et al.*, 2020; Schield *et al.*, 2020). Reduced recombination events should result in a loss of GC-biased gene conversion on former microchromosomes (Duret *et al.*, 2006; Ellegren, 2010), and with it a lower GC content on these regions. Conversely, the smaller regions that have translocated from larger chromosomes onto microchromosomes, are gaining GC content as a result of increased recombination and the GC-biased conversion that comes with it. As fused regions appear to be intermediate to conversed regions in GC content, it is plausible that these regions are still moving toward a new equilibrium. The more perplexing issue is the seemingly genome-wide loss of GC-content. Fusions of chromosomes are themselves a genome-wide phenomenon: while not all fusions have resulted in a change of chromosome size categories, we would expect all fusions to have increased chromosome size. As such, genome-wide AT-GC disequilibrium may be driven in part by an overall reduction in recombination rate. More detailed studies on specific chromosomal rearrangements within Falconidae will be necessary to explore this hypothesis.

We likewise detect a bias towards inserted bases relative to deleted bases in both small indels and large structural variants. These findings are in keeping with previous reports that falcons have the lowest microdeletion rate observed in birds (Kapusta *et al.*, 2017), and may also explain the longer genes, intragenic spaces, and introns in falcons than typical of other birds (Zhang *et al.* 2014). As Biased-Gene-Conversion has been shown to favor inserted bases over deletions, this result is at odds with our findings of AT-biased mutations and suggests that other mechanisms may be important toward driving insertions (Leushkin and Bazykin, 2013); however, the accumulation of insertions on former microchromosomes irrespective of current state likely reflect higher ancestral recombination rates. While insertion-biased-gene-conversion has been previously reported to account for interspecific fixations, our findings demonstrate that it can be observed intraspecifically as well. Importantly, indels and large structural variants seem to comprise a genomic feature for which ancestral chromosome types seem to be more important than current chromosome type. As such, conserved sequence motifs or other factors that intrinsically separate microchromosomes and macrochromosomes may remain on these fused regions and continue to affect structural variation (Perry *et al.*, 2020).

Analysis of TEs and chromosomal type confirms that repetitive elements generally tend to accumulate on larger chromosomes, as previously proposed (International Chicken Genome Sequencing Consortium, 2004; Boissinot *et al.*, 2019). Interestingly, regions of microchromosomes that have fused to larger chromosomes show intermediate levels of bases annotated as originated from repetitive elements. Given multilateral evidence for a dearth of repeat activity in falcons, it is probable that the intermediate levels of bases originating among repetitive elements is the result of reduced efficiency at removal of older repetitive bases on larger chromosomes. This finding is again consistent with lower recombination rate, as ectopic recombination typically drives removal of repetitive elements (Frahry *et al.*, 2015; Ji and DeWoody, 2016; Jedlicka *et al.*, 2020). A lack of repeat activity may itself be considered an additional form of genomic instability. The extent to which falcons have experienced inactivation of repeats requires further study. Repeat activation is often associated with speciation events (Jurka *et al.*, 2011), but the falcons that we sampled have undergone rapid radiations without any evidence for enhanced repeat mobilization. It is tempting to speculate on a special role for the reduction of repeat activity in falcon evolution, given the links between repeat content, radiations, recombination, and chromosome stability and the peculiar nature of falcons in all these areas.

### Conclusions

Taken together our findings provide evidence that falcon genomes have undergone a dramatic reduction in recombination rate that has thrown them out of AT-GC equilibrium and favored the accumulation of AT content. We link these findings in part to the substantial number of chromosomal fusions and microchromosome loss in falcons. While differences between microchromosomes and macrochromosomes have been well-characterized in the past (Ellgren, 2010; Perry *et al.*, 2020), little is known as to whether these differences are the result of chromosome size or sequence motifs. Our findings suggest that differences in base composition, recombination, and TE content between microchromosomes and larger chromosomes are largely products of chromosome size, and subject to disequilibrium following size changes. Conversely, structural variant and indel formation may be driven in part by factors other than size. Ultimately, our results show that falcon genomes are undergoing similar genomic changes to mammals following similar (albeit) lesser microchromosome loss events, and support a strong role for biased-gene-conversion in both maintaining and driving genomic equilibrium and disequilibrium.

## Methods

### DNA Extraction and Sequencing

Blood samples were obtained from female falcons of known ancestry during the course of normal veterinary care (Table 1). Samples were collected in accordance with IACUC protocol FS 18-0001. High-molecular weight DNA was extracted from samples using the Qiagen MagAttract Extraction kit (Hilden, Germany) as per manufacturer’s instructions, with an additional tissue lysis step to improve yield. DNA fragment size was verified to be approximately 50kb using pulse-field gel electrophoresis on a Bio-Rad Laboratories CHEF Mapper XA System (Hercules, California, USA). Libraries were prepared using the 10X Chromium Genome High Throughput (HT) Gel and Bead Kit Version 2 (Pleasanton, California, USA), barcoded with the Chromium i7 Multiplex kit, and sequenced on an Illumina Novaseq6000 using the NovaSeq6000 S4 300 Cycle Reagent Kit (San Diego, California, USA). Sequence quality was assessed using FastQC Version v0.11.8 (Andrews, 2017), and sequences were assessed for potential GC bias and contamination with Kat Version 2.3.1 (Mapleson, 2017).

### Genomic Assembly

Sequences were assembled using the Supernova assembly software for *de novo* assembly of linked-reads (Weisenfeld *et al.*, 2017). Sequences were not trimmed beforehand in accordance with the instructions of the software provider. Supernova provides several FASTA output options. Genomes were output using the “Pseudohaplotype 2” option to produce 2 phased haplotypes for each genome. Downstream analyses either used both sets of phased haplotypes to perform analyses on each diploid genome, or used single representative scaffolds from each phased haplotype to analyze haploid genomes. Scaffolds smaller than 16kb were discarded from all assemblies.

Assembly completeness was assessed using the BUSCO version 5.0 and the “aves_odb10” database and Augustus 3.4 to search for conserved avian single-copy orthologs (Simão *et al.*, 2015). BUSCO was run using an initial chicken model for protein structure and supplemented with a new search using a genome specific AUGUSTUS model based on discovered genes (option “long”).

Presence of Z and W sex chromosomes were identified by searching for diverged sex chromosome specific Z (CHD-Z) and W (CHD-W) Chromosome Helicase genes (Fridolfsson *et al.*, 1999) with HMMER (Wheeler and Eddy, 2013). In brief, HMMER profiles for both genes were created by downloading all versions of these genes from birds between 200 and 2200 base-pairs in length from NCBI Genbank (Benson *et al.*, 2012) using “esearch” from the Entrez tool utilities (Kans, 2017) on February 7, 2020 with the following specific search parameters: “txid8782[Organism:exp] AND biomol_genomic[PROP] AND (“200”[SLEN] : “2200”[SLEN]) AND chd-z[All Fields] OR chd-w[All Fields]”

These were aligned using a MAFFT version 7.407 (Katoh and Standley, 2013) iterative global alignment and used to construct a HMMER profile which was used with nHMMER to search for copies of these genes at an e-value threshold of 0.000001. Results were manually confirmed as CHD-Z and CHD-W with NCBI BLAST searches.

Mitogenomes were created by aligning raw reads to consensus genomes of archival mitogenomes. In brief, all mitogenomes for peregrine falcons, *Hierofalco*, and the common kestrel were downloaded from NCBI Genbank using E-utilities with the following respective search parameters: “txid8954[Organism] mitochondrion[filter] AND (“16000”[SLEN] : “20000”[SLEN])” “(txid345155[Organism] OR txid120794[Organism] OR txid345164[Organism]) mitochondrion[filter] AND (“16000”[SLEN] : “20000”[SLEN])” “txid100819[Organism] mitochondrion[filter] AND (“16000”[SLEN] : “20000”[SLEN])”

Archival mitogenomes were aligned using a Mafft local-pairs iterative alignment and constructed into a HMMER database. Archival mitogenomes were then aligned back to these preliminary consensus genomes with HMMER to determine common starting points. Mitogenomes were reoriented to the same starting point, realigned with a mafft local-pairs iterative alignment and final archival consensus genomes were created. Long Ranger was used to align raw reads back to the appropriate archival consensus mitogenome (i.e. Peregrine, *Hierofalco*, Common Kestrel) for each sequenced genome with the requirement that reference alleles have a minimum depth 3X higher than the reference allele.

### Variant Calling

Variant calling was performed using several methods. First, 10X Genomics Long Ranger (Marks *et al.*, 2019) was used to align raw demultiplexed reads from each sample back to their own haploid assemblies. In accordance with the proprietor’s instructions, GATK (version 3.5) was used as a variant caller with all 10X quality filters enabled. These filters excluded variants with: total Pfred quality <30; Pfred Quality of < 15 for any heterozygous allele; an allele fraction less than 15%; inconsistent phasing (based on linked reads); >3bp unphased homopolymer insertions (based on linked reads); and, poorly aligned variants calls of whole molecule relative to the scaffold (“10X_RESCUED_MOLECULE_HIGH_DIVERSITY”). While this approach worked well for calling variants within each genome, it was insufficient for alignments across falcon species due to higher sequence divergence. As such, MUMmer version 3.2.6 (Kurtz *et al.*, 2004) was used to align whole diploid genomes from each assembly to the haploid common kestrel genome. To facilitate alignments between species, MUMmer alignments used the NUCmer algorithm with a minimum alignment length of 100 bp and a maximum gap of 500 bp within a single alignment, in accordance with manual guidelines for “fairly similar sequences.” The common kestrel genome was chosen as a common genome for alignment due to its equal evolutionary distance from all other genomes, thereby avoiding biases in variants calls resulting from better alignments between more closely related species. Custom scripts were used to convert MUMmer alignments of small variants into vcfs and to annotate larger variants into bed files. To validate this approach, phased haplotypes of each diploid genome were aligned back to one another using MUMmer and the results were compared to the results of the Longranger output. Target haploid genomes for all alignment approaches had scaffolds smaller than 100 KB removed to better facilitate large-structural variant analysis. Large structural variants were annotated from MUMmer alignments to the common kestrel and between alternative pseudohaplotypes of the same genomes using a custom script. Large insertions and deletions were defined as a 50bp or greater breaks in alignment on one sequence, without any break in alignment on the other, with large deletions demonstrating the break on the reference scaffold and large insertions demonstrating the break on the aligned scaffold. Large inversions were defined as 50bp or greater reverses in direction of alignment between scaffolds, relative to the majority rule directionality of alignments up to that point between the same scaffolds. Large insertions, deletions, and inversions, were all filtered to exclude any matches to duplicated regions. However, segmental deletions and duplications were defined as 50bp or greater matches in alignments to single proximally aligned positions: segmental deletions were annotated when a tandem alignment of the reference genome matched a single upstream position of the aligned genome; segmental duplications were annotated where tandem regions of the aligned genome matched a single proximal region of the reference genome.

Small insertions and deletions are insertions and deletions of less than 50bp relative to the reference genome after normalization. Small insertions and deletions were left aligned and normalized prior to any analysis. All comparisons and processing of variants were performed using Samtools and Bedtools (Li *et al*, 2009; Quinlan and Hall, 2010; Li, 2011).

### Transcriptome and Gene Annotation

Samples were collected in RNA*later*® (Thermo Fisher Scientific: Waltham, Massachusetts, USA) from a female gyrfalcon immediately following clinical euthanasia for amyloidosis of the liver and frozen at −80°C until processing. In total, samples were taken from 22 distinct tissues: the liver, pancreas, chest muscle, stomach, hyperpallium (brain), cerebellum (brain), thyroid, gizzard, heart, kidney, lung, spleen, and at 10cm intervals in the intestine (with 10 intestinal samples total). All samples were extracted using the Qiagen RNA extraction kit, along with two negative controls—one taken from the beginning and one at the end of all extraction steps. Extracted RNA was normalized to 10ng in 50µl for library preparation. Library preparation was performed with the NEB Ultra II RNA Library Prep kit and sequenced in multiplex on a NovaSeq 2×150 flowcell.

Reads were used to produce a *de novo* transcriptome. In brief, reads were mapped to the Canadian gyrfalcon reference genome (Gyr-1) using HiSat2 (Kim *et al.*, 2019) and assembled into transcripts using StringTie (Pertea *et al.*, 2015). Transcript regions were identified with Transdecoder (Haas *et al.*, 2013) and the transcriptome was annotated using the Trinotate pipeline (Bryant, 2017). This transcriptome was then mapped to each genome to produce gene annotations using the “merge” option in stringtie. Gene ontology (GO Term) annotations were produced using BLAST2GO (Conessa *et al.*, 2005).

### Repeat Annotation

Repeats were annotated using RepeatModeler2 (Flynn *et al.*, 2020) and RepeatMasker (Tarailo-Graovac and Chen, 2009). Repeatmasker2 was run independently on each diploid falcon genome using rmBLAST as a search engine and the LTR structural modeling enable. The default max sample size of 243,000,000 was used for a total number of 403,000,000 bases sampled across all rounds; this had the effect of sampling ∼16.8% of each haploid falcon assembly. In total, 2,643 repeat families were detected across all eight diploid assemblies. These were filtered using Usearch “sortbysize” to remove all repeat families with less than 20 copies found in any given genome, leaving 1503 putative repeat families. These were pooled, sorted by length and clustered into merged consensus (“consout”) sequences at 80% (subfamily level) with USEARCH (Edgar, 2010) using the “cluster_fast”, resulting in 955 unique clusters. These were further filtered to remove any subfamilies that had not been reported in at least 40 copies across all genomes, leaving 238 consensus subfamilies. These were annotated according to the initial seed sequence used to produce the cluster with a custom bash script. Repeatmasker v4.1.2 was run on each diploid falcon genome using rmblast v2.11.0 as a search engine and the merged falcon consensus subfamily sequences as a custom library. The “-gccalc” option was enabled to correct for background GC levels in target sequences and the “-s” option for a more sensitive search. Alignments were saved and used to create a divergence curve using the included RepeatMasker utilities. To date past TE activity, the same repeat library and annotation methods were used to annotate additional avian genomes: the emu (GCA_016128335.1); the red jungle fowl (GCF_000002315.6); the red-legged seriema (GCA_009819825.1); the Swainson’s thrush (GCF_009819885.1); and, the kakapo (GCF_004027225.2).

We searched for active repeats comparing polymorphic structural variants against repeat annotations using Bedtools: a repeat was considered polymorphic if it made up at least half of an insertion structural variant and was contained entirely within an insertion structural variant. We searched for active repeats by searching for intact open reading frames. In brief, NCBI E-Utilities “esearch” was used to download intact repeats belonging to the LINE, LTR, and DNA repeat subfamilies according to the following search criteria:

#### LINES

“reverse transcriptase” AND complete[Title] AND (non-LTR[Title] OR LINE[Title] OR “Chicken Repeat 1”) AND (“1200”[SLEN] : “8000”[SLEN]) NOT “pseudogene”

#### LTRs

“reverse transcriptase” AND “complete”[Title] AND “LTR” NOT non-LTR AND (“1200”[SLEN] : “12000”[SLEN]) NOT “pseudogene”

#### DNA

“complete”[Title] AND Transposase AND (Harbinger OR hAT OR MULE) AND (“1200”[SLEN] : “10000”[SLEN]) NOT “pseudogene”.

In total, 66 LINE sequences, 122 LTR, and 106 DNA sequences were acquired. Each of these libraries was aligned using Mafft l-insi iterative local-pairs alignments with up to 1,000 iterations, and constructed into HMMER databases, which were used to extract repeat ORFs with nhmmer. We considered repeat ORFs to be potentially active if they exceeded minimum length requirements for active ORFs of their subclass based on reports from NCBI Genbank: 2,000 bp for LINEs; 2200 bp for LTRs; 1,200 bp for DNA transposons.

### Isochore Window Annotation

Genomes were divided into non-overlapping 100 KB windows and non-overlapping 1MB windows. Base composition, CpG sites, variants, and repeats were assessed for each window (Costantini and Musto, 2017). Windows were assigned as belonging to ancestral chromosomal states based on their alignments to chromosome scale NCBI Genbank/RefSeq genomes for the Swainson’s thrush (*Catharus ustulatus*: GCF_009819885.1), kakapo (*Strigops habroptila*: GCF_004027225.2), and red-legged seriema (*Cariama cristata*: GCA_009819825.1). These birds were chosen to sample across all major clades related to falcons: the parrot and song bird clades of Eufalconimorphae (Suh *et al.*, 2011), and the outgroup to this clade (Prum *et al.*, 2015), respectively. They were also intended to sample birds across a wide range of chromosomal configuration: Swainson’s thrush has a typical avian chromosome count with 2N=80 annotated autosomes. Kakapos have a reduced chromosome count with 2N=46 annotated autosomes; and, the red-legged seriema has a more fractured genome with 2N=100 annotated autosomes. Whole genomes from each falcon were aligned to each of these genomes using MUMmer with the NUCmer algorithm, a minimum alignment length of 100 bp and a maximum gap of 500 bp within a single alignment. Windows were assigned as ancestrally belonging to: microchromosomes if they aligned exclusively to chromosomes of less than 20MB across all three non-falcon genomes, intermediate chromosomes if aligned exclusively to chromosomes greater than 20MB and less than 40MB across all three non-falcon genomes, and macrochromosomes if they aligned exclusively to chromosomes greater than 40MB across all three falcon genomes. “Large chromosomes” were defined as any chromosome greater than 20MB. Window’s aligning to chromosomes of different size categories within or across the three non-falcon genomes were classified as belonging to an ambiguous ancestral state and removed from analyses relating to ancestral state. The current state of windows as belonging to microchromosomes, intermediate chromosomes and macrochromosomes was annotated in an identical manner, except with alignment to chromosome scale assemblies of the gyrfalcon (GCF_015220075.1) and peregrine falcon (GCA_009819825.1) found on NCBI RefSeq and Genbank, respectively. Windows belonging to sex chromosomes were also annotated using the same approach; as the peregrine assembly was obtained from a male it contained only a Z Chromosome, and windows belonging to the W Chromosome could not be annotated in this way and were excluded from windows-based analysis. All statistical analyses relating to base-composition were performed on 100 KB windows in accordance with typical isochore size and standard practices for isochores annotation (Costantini and Musto, 2017). Base composition and CpG sites were determined using the program SeqTK (Shen *et al.*, 2016).

Graphs were made for falcon and other amniotes using the following archival sequences: the kakapo (GCF_004027225.2); the Swainson’s thrush (GCF_009819885.1); red-legged seriema (GCA_009819825.1); the saltwater crocodile (*Crocodylus porosus*: GCF_001723895.1); the green sea turtle (*Chelonia mydas*: GCF_015237465.1); sand lizard (*Lacerta agilis*: GCF_009819535.1); and a human (*Homo sapiens sapiens*: GCF_000001405.39).

### Nuclear Mitochondrial DNA (NUMT) Annotation

NUMTs were annotated using HMMER. In brief, all mitochondrial genomes from Falconidae on NCBI were downloaded on October 4, 2021, using NCBI Entrez tools “esearch” with the search terms: “txid8949[Organism] mitochondrion[filter] AND (“16000”[SLEN] : “20000”[SLEN])”.

Several additional mitogenomes were downloaded in the same manner to assure taxon-sampling across Aves: *Catharus ustulatus* (CM020378); *Cacatua alba* (MT920475); *Picus canis* (NC_045372.1); *Strix occidentalis* (MF431746); *Accipiter gentilis* (AP010797); *Gavia stellata* (AY293618); *Balearica regulorum* (FJ769841); *Columba livia* (KP319029); *Gallus gallus gallus* (AP003322); *Struthio camelanus* (AP003322).

Mitogenomes were aligned using the Mafft local-pairs iterative alignment with up to 1,000 iterations, and aligned into a HMMER database. Falconidae mitogenomes were realigned to the HMMER database with nhmmer and then reoriented to the same start position. Reoriented mitogenome sequences were then concatenated with themselves to ensure that all regions were fully present without a break point. These concatenated and reoriented mitogenomes were then realigned with a Mafft local-pairs iterative alignment with up to 1,000 iterations, and compiled into a HMMER database. This database was used to search the full diploid genomes of all falcons with nhmmer using an e-value of 10^-6^. Synteny analysis was performed by converting the locations of NUMTs in all falcons to locations in the kestrel genome based on MUMmer alignments using a custom script; locations were then compared with Bedtools. Extracted NUMTs were clustered at 80% using USEARCH and trees were constructed for each cluster using RAxML (Stamatakis, 2014) with a GTR model, including gamma-distributed rate. Trees were built using the best of 5 guide trees, and bootstrapped 20 times. Analyses were conducted with NUMTs that occurred in only a single cluster.

### Phylogenetic Tree Construction

SNV calls from MUMmer alignments to the haploid kestrel genome were used to create aligned full genome consensus sequences for each genome using Bcftools “Consensus” in Samtools. These were converted to gvcfs using a custom script and filtered against conserved sites to extract aligned vcfs of variable sites in each genome. These vcfs were converted into Fasta alignments with a custom script and Fasta alignments were converted to Phylip alignments with the program Readseq (Gilbert, 2003).

A maximum-likelihood tree was constructed in RAxML with all variable sites using a GTR model with gamma-distributed rate variation between sites. The common kestrel was specified as an outgroup. Bootstraps were performed in 1000 replicates and drawn onto the best tree to assess branch support. The tree was scaled to time and dated using the program R8S (Sanderson, 2003). The split between the common kestrel and other falcons was used as a calibration point. This split was specified as 7.246 million years ago, corresponding to the Tortonian-Messinian junction in the upper Miocence based on the fossil record of the earliest diverged member of the old-world kestrels (Boev, 2011) and in accordance with previous molecular-clock-based estimates of this split (Fuchs *et al.*, 2015). Divergence times were estimated using the penalized-likelihood approach (Sanderson, 2002) and the truncated newton algorithm.

### Pairwise Sequential Markovian Coalescent

Analysis of falcon demography was performed using a Pairwise Sequential Markovian Coalescent (PSMC) analysis with the program PSMC (Li and Durbin, 2011). The model was parametrized with a maximum coalescent time of 12 N_e_ generations, theta/rho of 2, and modelled across 100 time-intervals with separate consensus estimates for the first and last eight intervals and 28 three-interval segments between these. Theta/rho was estimated based on the per-generation mutation rate, mu, of 4.6 x 10E-9 estimated from the collared flycatcher (Smeds *et al.*, 2016), and the per-generation recombination rate of 0.0014 in the zebra finch (Singhal *et al.*, 2015). All PSMCs were bootstrapped 100 times.

Generation times were estimated using the equation: [1] g = a + [s ⁄ (1 - s)], where “a” is the age of sexual maturity and “s” is the expected adult survival rate (Zhan *et al.*, 2013). This provides a generation time of 3.13 years for the common kestrel (Dijkstra *et al.*, 1990), 6 years for the peregrine falcons (Court *et al.*, 1989; Kauffman *et al.*, 2003), 6.56 years for the saker falcon (Wink *et al.*, 1999; Kenward *et al.*, 2007), and 12 years for the gyrfalcons (Booms. 2010). Reliable estimates of survival could not be found for lanner falcons, and a generation time of six years was used.

### Statistical Analyses

All statistical analyses were run in R Studio 1.1,1456 (R Studio, 2016) using R version 3.5.1 “Feather Spray” (R Core Team, 2018) Isochore analysis was conducted using linear models. Models comparing current and ancestral chromosomal states were chosen using stepwise selection based on AIC values with both forward and backward selection enabled. Where GC content was included in the model, it was regressed against the response variable to remove its effect and subsequent analyses were performed on residuals. As count data such as those used in this analysis typically has variance in proportion to mean, log_2_(x+1) transformations were performed on all datasets to normalize residuals. Anderson-Darling tests were used to assess normality of residuals and confirm the efficacy of transformations, which resulted in an approximate 10-50X decrease in the size of scale of the Anderson-Darling statistic (Anderson and Darling, 1954). Anderson-Darling tests were performed using the “nortest” package. Comparison of levels of between chromosomal states were performed using Tukey’s “Honest Significant Difference Method.” We use a false-discovery-rate corrected alpha value of 0.0032 in accordance with a Dunn–Šidák correction of an initial alpha of 0.05 for our 16 independent statistical tests.

## Data Availability Statement

The genome sequences have been deposited at GenBank under BioProject accession PRJNA 794321.

## Acknowledgements

This work was supported by funding from the Center for Genomics and Systems Biology at New York University Abu Dhabi. The NYUAD Dalma and NYU Prince and Greene high-performance-computing clusters were used to produce this work. Sequencing work was performed by Marc Arnoux of the Center for Genomics and Systems Biology at NYUAD. Bioinformatics work was performed by Dr. Nizar Drou of the Center for Genomics and Systems Biology at NYUAD. We thank DR. Sebastian Kirchhof for his comments on an earlier version of the manuscript. We would also like to thank Dr. Peter McKinney of Al Aseefa Falcon Hospital and Mohammed Al Kamda of Al Kamda Falcons for consulting on this project.

